# Joint automatic metabolite identification and quantification of a set of ^1^H NMR spectra

**DOI:** 10.1101/2020.10.08.331090

**Authors:** Gaëlle Lefort, Laurence Liaubet, Nathalie Marty-Gasset, Cécile Canlet, Nathalie Vialaneix, Rémi Servien

## Abstract

Metabolomics is a promising approach to characterize phenotypes or to identify biomarkers. It is also easily accessible through NMR, which can provide a comprehensive understanding of the metabolome of any living organisms. However, the analysis of ^1^H NMR spectrum remains difficult, mainly due to the different problems encountered to perform automatic identification and quantification of metabolites in a reproducible way. In addition, methods that perform automatic identification and quantification of metabolites often do it for one given complex mixture spectrum. Hence, when a set of complex mixture spectra coming from the same experiment has to be processed, the approach is simply repeated independently for every spectrum, despite their resemblance. Here, we present a new method that is the first to identify and quantify metabolites by integrating information coming from several complex spectra of the same experiment. The performances of this new method are then evaluated on both simulated and real datasets. The results show an improvement in the metabolite identification and in the accuracy of metabolite quantifications, especially when the concentration is low. This joint procedure is available in version 2.0 of **ASICS** package.

## 1 Introduction

Among omics, metabolomics are promising to identify potential biomarkers as they are relatively close to the final phenotype and because of their moderate cost [6]. Nuclear Magnetic Resonance (NMR) allows to obtain metabolomic profiles from easy-to-obtain fluids (*e.g*., plasma, serum or urine), and NMR spectrometers produce a spectrum from a sample of one of these fluids. We will term such a spectrum a “complex spectrum” as it provides a profile of the quantification of all the metabolites contained in the sample [11]. However, the quantification is not direct: the complex spectrum is made of several peaks, where one peak can correspond to several metabolites, and one metabolite is described by one or several peaks –the quantity of the metabolite in the sample varying proportionally to the area under its peaks.

The classical approach to analyze such spectra consists in cutting each spectrum in small intervals, called buckets, and in computing the area under the spectrum of each bucket to perform statistical analyses [1, 22]. Since buckets are not directly connected to metabolites, this approach requires that NMR experts identify the metabolites from the buckets that are found relevant by the statistical analyses for a given biological question. This identification step is tedious, time consuming, expert dependent and, by consequence, not reproducible [18]. It also leads to a serious loss of information since the identification of metabolites is restricted to the metabolites that correspond to extracted buckets [5].

To ease the use of NMR data, we developed a method, **ASICS**, which allows to automatically identify and quantify metabolites in NMR complex spectra [10, 16] (R Bioconductor package at https://bioconductor.org/packages/ASICS/). This method is based on a library of pure spectra (*i.e*., spectra obtained from a single metabolite) that is used as a reference to fit a reconstruction model. This method has been evaluated in Lefort *et al*. [10], where quantifications of metabolites were performed on urine of diabetic patients and on plasma of pig fetuses and were compared to a manual identification and quantification performed on a few targeted metabolites. Overall, the comparison showed that the automatic quantification provided results similar to the expert manual processing but in a much shorter amount of time and with an easily reproducible procedure. It also showed that **ASICS** had a much better sensitivity / specificity trade-off than other automatic identification methods such as **batman** [8] or Bayesil [13] and improved quantification compared to targeted automatic quantification methods like **rDolphin** [3] or Autofit [19].

However, we also showed that quantifications of less concentrated metabolites were of poorer quality, as is often the case in automatic methods, because these metabolites are hard to distinguish from the noise level. This problem may stem from quantification itself but also from preprocessing steps that come prior to the quantification and aim at improving the quality of the analyzed complex spectrum. Among critical preprocessings, one of them aims at aligning every pure spectrum of the reference library on the analyzed complex mixture. **ASICS** uses its own alignment, inspired by the alignment implemented in **speaq** [2], but NMR tools include methods that were also designed to perform spectrum alignment, like icoshift [14] or **speaq** [2]. However, whatever the identification and quantification tools, they are all designed to process the complex spectra one by one, independently, which is under-efficient when these come from the same experiment in closed conditions and thus share some similarities with one another.

Here, we present a new method to align pure spectra with the complex spectra of a sample of interest and to estimate quantifications that integrate information obtained from several complex spectra of the same experiment. The joint alignment is performed by automatically calibrating one of the parameters of the alignment algorithm. The joint quantification uses the joint alignment and is based on the use of a multivariate regression model incorporating a group sparse penalty. Both approaches are evaluated on simulated spectra (for which a ground truth is available) and on a real dataset of newborn piglet plasma and lead to improved identification and quantification, especially for lowly concentrated metabolites. This joint procedure is available in version 2.0 of **ASICS** package.

## 2 Methods and tools

### 2.1 General overview of the quantification strategy

Automatic identification and quantification of metabolites in a complex spectrum, **f**, is performed using a reference library of *p* pure spectra, (**g**_*j*_)_*j*=1,…,*p*_ (*e.g*., spectra obtained from a single metabolite) [10, 16]. The method then fits a model where the complex spectrum is decomposed into a linear combination of pure spectra in which the estimated coefficients divided by the number of proton *u*_*j*_ of the metabolite *j*, (***β***_*j*_)_*j*_*/*(*u*_*j*_)_*j*_, correspond to the quantification of the corresponding metabolites j ∈ {1, …, *p*}. To obtain valid quantifications, the coefficients (***β***_*j*_)_*j*_ are thus additionally constrained to be positive or null, which leads to the following model:

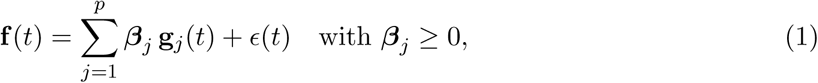

where **f** (*t*) and (**g**_*j*_(*t*))_*j*=1, …,*p*_ respectively correspond to the complex spectrum to quantify at chemical shift *t* (in ppm) and to the *jth* spectrum in the reference library also at chemical shift *t*. The noise *ϵ*(*t*) is assumed to be structured such that *ϵ*(*t*) **╨** *ϵ*(*t′*) for *t* ≠ *t′* and includes both an additive noise *ϵ*_2_(*t*) and a multiplicative noise *ϵ*_1_(*t*) such that: 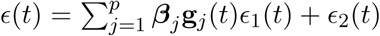 where 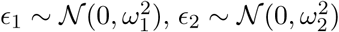 and *ω*_1_, *ω*_2_ are user-defined values (`mult.noise` and `add.noise` respectively in **ASICS** R package).

However, the model is fitted only after a number of preprocessing steps have been performed as illustrated in Fig. 1 (see Lefort *et al*. [10] for further details): a **library cleaning step** selects a limited number of relevant pure spectra in the reference library to be used in model (1) in order to improve its fit. Then, **two alignment steps** are performed to align the peaks of every selected pure spectrum, **g**_*j*_, to the peaks of the complex spectrum **f**. These steps are necessary to correct peak shifts or distortions (expansion or narrowing) due to technical variations during the acquisition process (*e.g*., pH or temperature). A global shift, *s*_*j*_, is first estimated individually for every pure spectrum **g**_*j*_ and a refinement of this shift is then performed for every peak in **g**_*j*_ to estimate additional local shifts.

**Figure 1:**
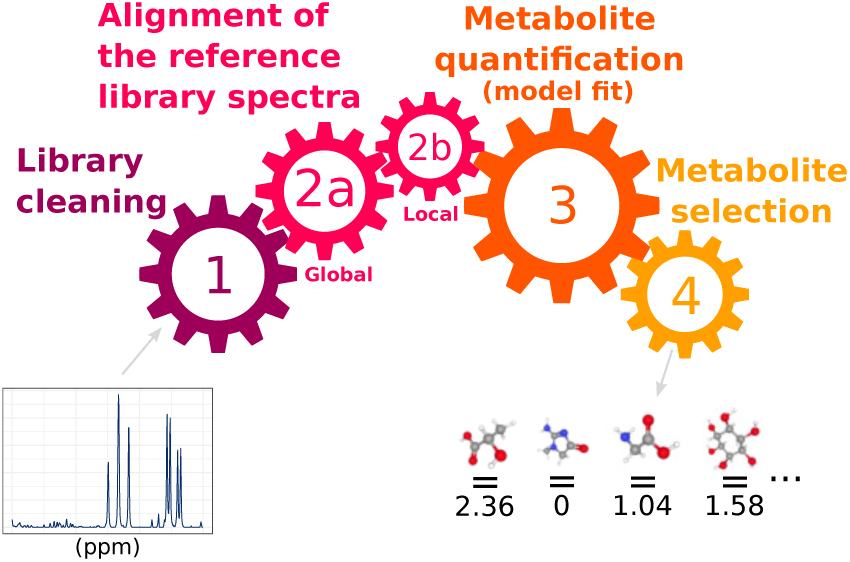
Steps of the metabolite quantification of NMR spectra.

In addition, a postprocessing step is performed after the model of Equation (1) has been fitted. It aims at controlling the number of falsely selected metabolites. A **multiple testing selection procedure** based on FamilyWise Error Rate (FWER) is performed and consists in computing a threshold, *ν*_*j*_, for each metabolite, which depends on all the estimated parameters (***β***_*j*′_)_*j*′_=1,…,*p* and then in setting to 0 estimates (*i.e*., quantifications) such that {***β***_*j*_ ≤ *ν*_*j*_}.

The following two paragraphs will describe in more detail the two alignments steps, for which a joint version is proposed in this article. These alignments cannot be performed with usual NMR alignment methods because peaks are much more rare in pure spectra than in complex spectra and are thus harder to precisely bound (in complex spectra, a peak is naturally bounded by its neighbor peaks). Technical drifts are also generally larger because pure spectra usually cannot be acquired in the same batch of experiments. The solution consists in first obtaining a global shift *s*_*j*_ by optimizing:

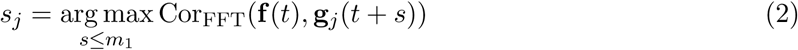

where Cor_FFT_ is the the fast Fourier transform (FFT) cross-correlation [20] between the complex mixture **f** and a set of pure spectra **g**_*j*_ shifted by *s* ≤ *m*_1_ with *m*_1_ a maximum shift defined by the user.

Then, each peak of the pure spectrum is independently aligned on the complex spectrum **f** locally, using a warping function that is constrained with a local maximum shift, *m*_2_ = *m*_1_*/*5. These two alignment steps result in an aligned reference library corresponding to the complex spectrum **f** whose quality is thus strongly conditioned on the user-defined parameter *m*_1_.

When the quantification is performed on *n* complex spectra, (**f**_*i*_)_*i*=1,…,*n*_, from the same experiments, a naive approach would be to perform all these steps *independently* for each complex spectrum. This would result in *n* different selections of the metabolites to be included in the model (library cleaning step) and in *n* aligned reference libraries. These aligned reference libraries all depend on a unique maximum allowed shift, *m*_1_, defined by the user and that generates global shifts, (*s*_*ij*_)_*j*_, and local shifts specific to the corresponding complex spectrum **f**_*i*_. In addition, in Equation (1), the error term *ϵ*_*i*_(*t*) and the estimated coefficients (***β***_*ij*_)_*i*_ would all depend on the complex spectrum under study, independently from each other, as well as the thresholds, (*ν*_*ij*_)_*i*_ that control the FWER.

However, complex spectra from the same experiment share some common traits. It is thus expected that using joint steps, in which cleaning, alignment and quantification are somehow “constrained” to share similarities between all complex spectra of a same experiment or of a same condition within an experiment, has the potential to improve the overall quality of metabolite identification and quantification. In the next two sections, we describe two procedures for joint reference library alignment and joint metabolite quantification, respectively. Note that these two procedures are not meant to be used together (Fig. 2): joint alignment aims at providing *n* aligned reference libraries for which the maximum shift allowed, *m*_1_, is optimally and automatically tuned for information coming from all spectra rather than being user defined. This refined joint alignment has the potential to improve quantification of the metabolites when the model (1) is fitted independently for each complex spectrum (as illustrated in the Section “Results and discussion”).

**Figure 2:**
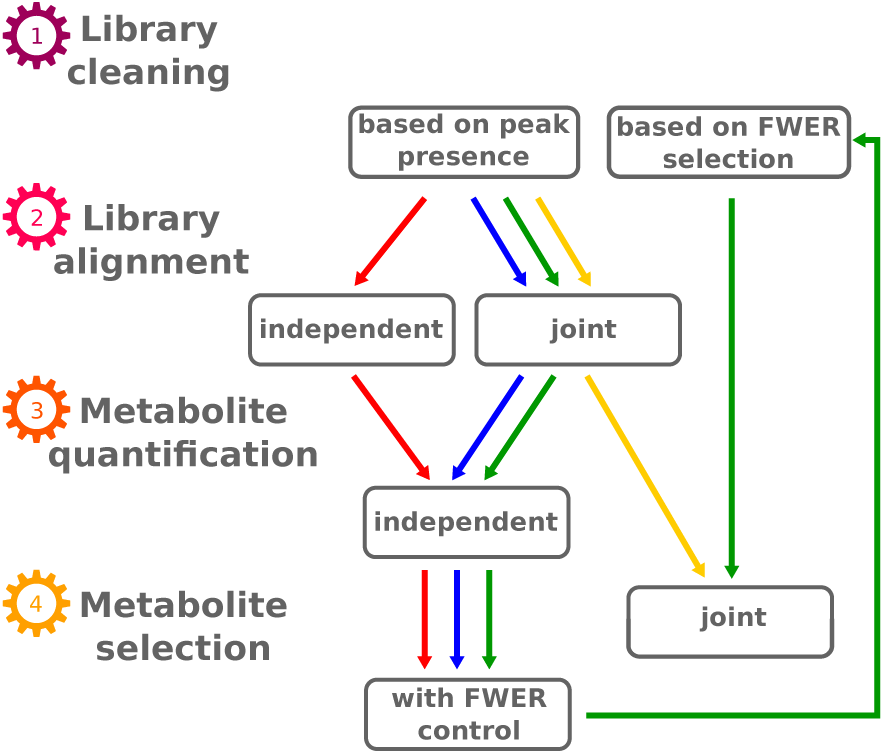
Four different scenarios for automatic metabolite quantification (red, blue, green and yellow arrows).

On the other hand, the joint quantification is a globally joint procedure that uses an aligned library that is common to the *n* complex spectra. It thus includes its own alignment step, derived from the joint alignment procedure and called the “common alignment” step. Advantages and drawbacks of these two joint approaches, depending on the experiment characteristics and on the user’s expectations, are discussed in the Section “Results and discussion”.

### 2.2 Joint alignment of the reference library

In the previously described alignment steps, the reference library is aligned independently on all complex spectra **f**_*i*_ and all pure spectra in the reference library **g**_*j*_ but this alignment depends on a unique maximum allowed shift, *m*_1_, used for both the global and the local alignments. This parameter somehow represents the “typical maximum shift” expected for the experiment and it is critical to properly set the range of values that are maximized with the Corr_FFT_ measure as in Equation (2). Previous experiments have shown a rather important sensitivity to this parameter and also that its value would be better determined depending on a given pure spectrum, **g**_*j*_, because it presents high variations in relation with the range of the spectrum shifts.

The idea of the joint alignment of the reference library is therefore to automatically set a specific maximum allowed shift for each pure spectrum, *m*_1*j*_, using information obtained from all complex spectra. This method thus increases the number of maximum shifts from 1 to *p* and provides more flexibility to account for the difference between pure spectra, while being more adapted to the given set of complex spectra. It is summarized in Fig. 3 and the full method is given in Algorithm 1.

**Figure 3:**
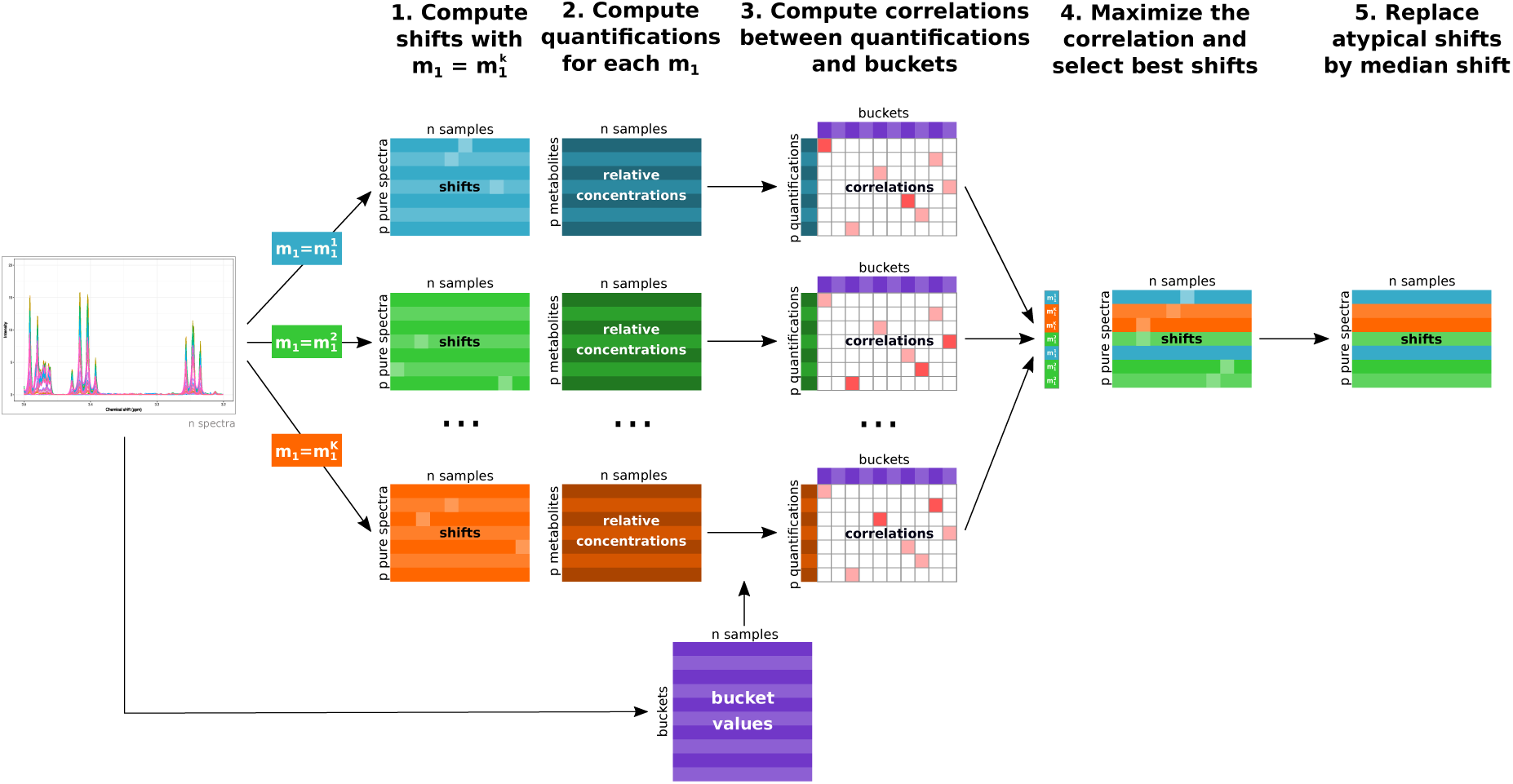
Overview of the different steps of the joint alignment of the reference library.

More precisely, for a given pure spectrum **g**_*j*_, *m*_1*j*_ is tuned by performing a rough quantification based on several maximum shift candidates (steps 3-4 of the algorithm) and by independently computing a measure of fitness between this estimated quantification and a bucket area for all complex spectra (step 5). Even if the bucket area is a poor estimate of the true metabolite quantification, having this information from several complex spectra allows to make it usable to compute a relevant quality measure of the alignment preprocessing (step 10) and thus of each maximum candidate shift. The “best” maximum shift is therefore finally selected from this quality measure (step 12).

Global and local alignments of every pure spectrum **g**_*j*_ are performed for all complex spectra (**f**_*i*_)_*i*_ using the estimated maximum shift *m*_1*j*_, and an additional joint post-processing step is then performed: the global alignment results in the computation of global shifts (*s*_*ij*_)_*i*_, all smaller than *m*_1*j*_. Outlier shifts (*s*_*ij*_ for which |*s*_*ij*_ *−*median(*s*_*i′j*_)_*i′*=1,…,*n*_| *>* 5*×*(*t*_2_*−t*_1_)) are thus further corrected and replaced by median(*s*_*i′j*_)_*i′*=1,…,*n, i*′≠*i*_.

### 2.3 Joint metabolite quantification using a multivariate Lasso

In the standard procedure where complex spectra are all processed independently from one another, the identification of metabolites present in a given complex spectrum **f**_*i*_ is performed by a postprocessing step performed after the model fit. This procedure uses thresholds, *ν*_*ij*_, based on FWER control, that are obtained independently for each complex mixture **f**_*i*_ and allows to decide whether the metabolite *j* should be selected or not. This approach allows to control the FWER of the metabolites in every complex spectrum **f**_*i*_ but can suffer from a lack of power. Since complex mixture spectra of a same experiment are expected to share a large fraction of common metabolites, the identification power of the procedure could be improved by using information from all spectra rather than performing the selection independently. In addition, in this independent approach, the quantification (model fit) and the identification (FWER control) are performed in two consecutive steps. The idea of the proposal described in this section is to address these two issues by designing a joint approach with a simultaneous identification and quantification that are based on all complex spectra at a time.

#### Algorithm 1 Joint alignment of the reference library.

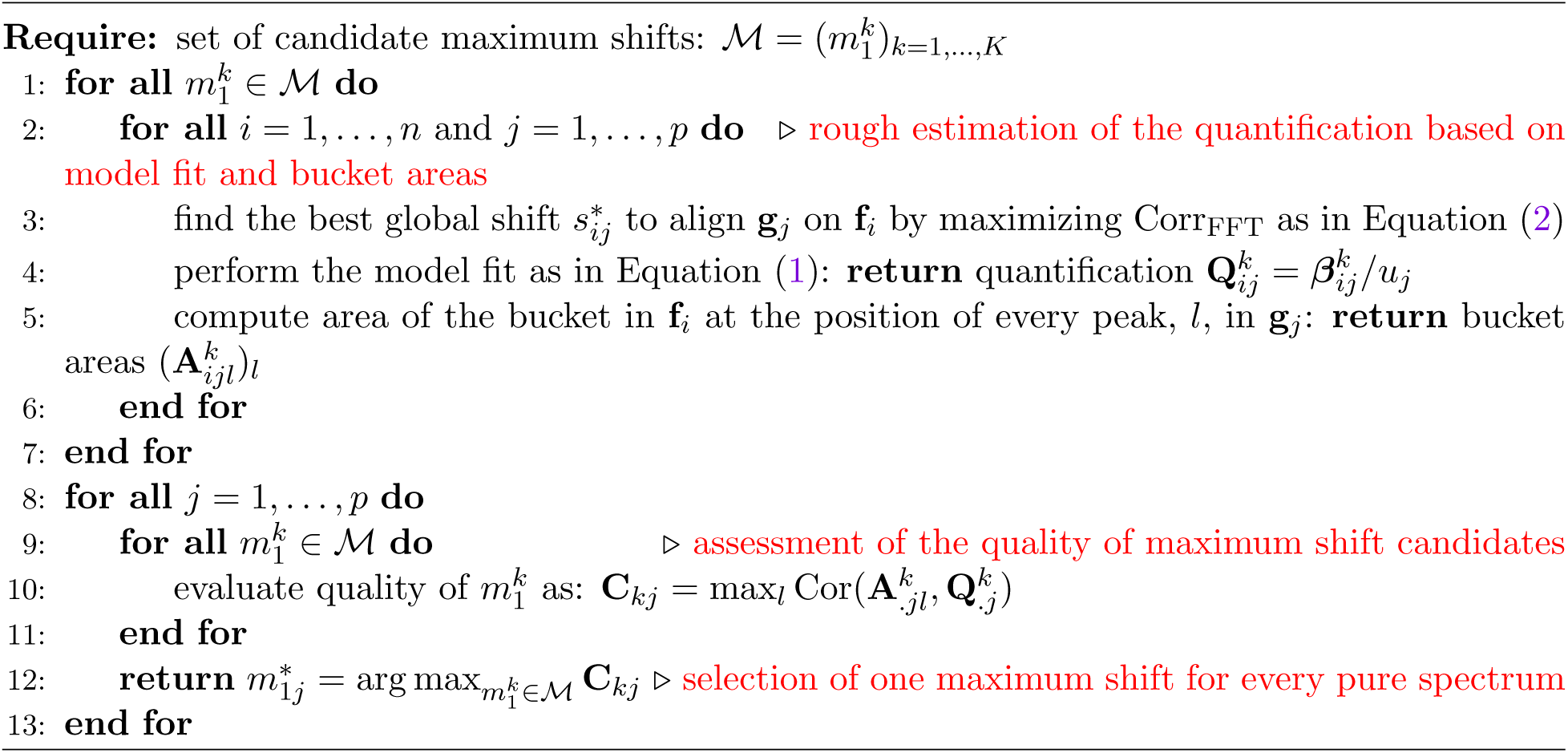

To do so, the idea is to fit a multi-response version of model (1), in which the simultaneously predicted values are **F**, the (*q × n*)-matrix of columnwise complex spectra (**f**_*i*_)_*i*=1,…,*n*_. This requires to obtain an aligned reference library common to all complex spectra, **G**, which is made of the (*q × p*)-matrix of columnwise aligned pure spectra (**g**_*j*_)_*j*=1,…,*p*_ and will serve as predictor of the multi-response version of model (1). In short, this common aligned reference library is based on the same preprocessing and postprocessing steps as the ones described in Fig. 1 that are aggregated using, for instance, a user defined ratio of common evidence within complex spectra, *r*_*c*_. It contains two cleaning steps, designed to reduce the size of the reference library, *p*, and global and local alignment steps. Technical details on how the common aligned library is obtained are provided in Algorithm S1 of Supplementary material.

The multi-response model is then based on a matrix version of the least square minimization problem used to solve model (1), which writes:

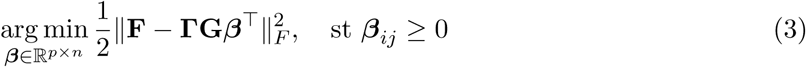

where **G** is the diagonal covariance matrix of the residuals and ‖· ‖_*F*_ is the Frobenius norm. In this version, the quantification of the metabolite associated to pure spectrum **g**_*j*_ in complex spectrum **f**_*i*_ are based on the estimated coefficient ***β***_*ij*_.

In addition, the use of a Lasso-type penalty to the square loss of Equation (3) is known to be efficient for selecting variables [17]. This type of penalty indeed enforces the sparsity of the solution of the minimization problem, *i.e*., the estimated coefficients (***β***_*ij*_)_*i,j*_ are forced toward 0, except for those most important for the prediction quality. In our case, a desirable property would be that all (***β***_*ij*_)_*i*=1,…,*n*_ are forced toward 0 simultaneously for a given *j, i.e*., that a given metabolite *j* is jointly identified or not identified for all samples. This can be performed by the use of a group-Lasso approach [21], that is based on the *ℓ*_1_-*ℓ*_2_ norm 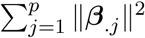, with ***β***_.*j*_ the vector of length *n*, (***β***_*ij*_)_*i*=1,…,*n*_.

Finally, the solved minimization problem is identical to the one implemented in the R package **glmnet** [7] and described in Simon *et al*. [15]:

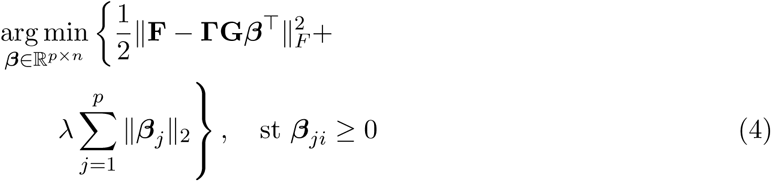

The parameter *λ >* 0 is used to control the trade-off between the adequation to the data (the error term computed with the Frobenius norm) and the model sparsity. It is usually tuned by cross-validation.

### 2.4 Implementation

Joint alignment and quantification are implemented in **ASICS** package version 2.0 (R Bioconductor package at https://bioconductor.org/packages/ASICS/). The user can define which approach to use (spectrum-dependent or joint alignment or quantification) by setting the following arguments:

`joint.alignment` to decide whether a joint alignment (if `joint.alignment=TRUE`) or an independent alignment (otherwise) is performed;

`quantif.method` to decide which type of quantification to perfom. The choices are either `“FWER”` (independent quantification for every complex spectrum), `“Lasso”` (not including “Cleaning step 2” for common library alignment) or `“both”` (including “Cleaning step 2” for common library alignment). The fit of model (4) is performed using the R package **glmnet** (version 3.0-2) and the regularization parameter, *λ*, is also tuned by the cross-validation procedure available in this package.

Note that if `quantif.method` is not set to `“FWER”`, the argument `joint.alignment` has no effect since the common alignment procedure of Algorithm S1 of Supplementary material is automatically performed;

`clean.threshold` to set *r*_*c*_ when a joint quantification is performed.

## 3 Experimental data and design

The joint alignment and joint quantification performances were assessed separately using two datasets: a simulated dataset was first used because of the ease to obtain a ground truth (true shift or true quantification) for performance quantification. A real dataset, in which some metabolites have been directly quantified using dosages, was also used to evaluate both aspects (but with no ground truth available for the shift, the alignment quality was evaluated indirectly by its impact on the quantification quality). Our approach was also compared with state-of-the-art alternatives freely available to perform alignment and/or quantification.

### 3.1 Simulated spectra

To assess the performances of joint alignment and joint quantification, we first simulated *n* spectra (**f**_*i*_)_*i*=1,…,*n*_ with metabolites in known concentrations, 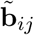, from some of the *p* pure spectra (**g**_*j*_)_*j*=1,…,*p*_ present in **ASICS** reference library. Parameters used to calibrate distributions for quantification simulations and shifts were obtained from previously analyzed real datasets and the precise steps of the simulations are described in Section S2.2 of the Supplementary material. They resulted in *n* = 100 simulated complex spectra, each composed of approximately *p*_*i*_ ∼ 95 pure spectra that correspond to metabolites in known concentration. The complex spectra were simulated in accordance with the model of Equation (1), as shown in Equation (S1) of the Supplementary material. The simulation process itself was repeated to obtain 100 such datasets.

### 3.2 Plasma spectra of newborn piglets

In addition, the performances were also assessed on newborn pig metabolome, obtained during the SuBPig project (funded by INRAE GISA 2018-2019). In this project, ^1^H NMR spectra were acquired on a Bruker Avance III HD NMR spectrometer (Bruker SA, Wissembourg, France) operating at 600.13 MHz for ^1^H resonance frequency from plasma of 97 Large White newborns collected on umbilical cord. NMR raw spectra are available in the Metabolights database [9]: MTBLS2137. The same samples were also used to obtained the concentrations of 27 targeted amino acids measured with an Ultra Performance Liquid Chromatography (UPLC). Details on the experimental protocol are available in Section S2.2 of the Supplementary material and basic statistic on amino acid dosages are provided in Table S1.

NMR spectra were quantified using **ASICS** with default parameters except for the threshold under which the signal is considered as noise that was set at 0.01 and the multiplicative and additive noise standard deviations that were set at 0.07 and 0.09 respectively. Noises were set at realistic values using fourteen technical replicates of a pool sample. The alanine peak (1.47–1.50 ppm) was used to set the multiplicative noise and the noisy area (9.4–10.5 ppm) was used to set the additive noise.

### 3.3 Evaluation of the joint alignment

The joint alignment procedure was compared to independent alignment as performed in **ASICS** and in two other tools designed for that purpose: icoshift [14] (version 3.0) and **speaq** [2] (version 2.6.1). All alignment methods were run for both datasets (simulated dataset and piglet plasma dataset) and, on the simulated dataset, 100 simulations of 100 complex spectra were performed to ensure the robustness of the results. In addition, assessment of the performance was not obtained identically for both datasets.

**For the simulated dataset**, a cosine similarity was computed for any metabolite *j* between the true (unknown) contribution of its given pure spectrum, **g**_*j*_, to the simulation of **f**_*i*_ (ground truth) and the result of the alignment of **g**_*j*_ on **f**_*i*_. For the sake of simplicity, this similarity was computed using the alignment obtained on a single reference complex spectrum, **f**_*i*\*_, that was the most similar (in terms of average cosine similarity) to all other complex spectra. This measure allowed to use the ground truth of the simulation to assess the quality of the alignment in a simple and efficient way.

In addition, the non-parametric Durbin test [4] (as implemented in the R package **PMCMR** [12]) was used to test the significance of the differences in cosine similarity between different alignment methods. The Durbin test allows to account for the pairing of metabolites across experiments and is also able to cope with the incompleteness of block design that is due to the fact that different metabolites are used to generate the reference complex spectrum **f**_*i*\*_ across experiments.

Once the reference library had been aligned, it was submitted to the **ASICS** independent quantification algorithm. The effect of the quality of the alignment on the quality of the identification and on the quantification was assessed. The metabolite identification quality was evaluated by comparing the identified metabolites with the metabolites truly used in the simulation. The significance of the difference in method sensitivity and specificity was assessed using Kruskal-Wallis test followed by the post-hoc Nemenyi test.

Finally, the metabolite quantification quality was evaluated by computing the correlation between the estimated metabolite quantification **b**_*j*_ and the ground truth metabolite quantification 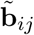 across *i* = 1, …, *n*. As for alignment quality, the significance of the differences between methods was tested using the Durbin test.

**For the piglet plasma dataset**, we did not know all metabolites that were truly present in the complex spectra so we could not perform the direct evaluation of the alignment quality, nor the evaluation through the quality of metabolite identification. However, we were able to assess the impact of the alignment on the quality of some metabolites’ quantification. This was done by computing correlations between estimated quantifications and UPLC concentrations, which are here considered as ground truth. The significance of the differences between methods was tested using the Durbin test followed by post-hoc Durbin tests.

### 3.4 Evaluation of the joint quantification

Different scenarios of the joint quantification method were evaluated: joint quantification with a single cleaning step (in yellow in Fig. 2), joint quantification with a second cleaning step (in green in Fig. 2), for which several values of the ratio of common evidence (*r*_*c*_) were tested: *r*_*c*_ *∈* {1%, 10%, 50%}. This joint quantification procedure was compared with quantifications obtained with **ASICS** independent quantification (in blue in Fig. 2). On the piglet plasma dataset, we also compared the results with another quantification method, performed independently on each complex mixture spectrum: the one implemented in the R package **rDolphin** [3] (which was the alternative quantification method which performed best among those tested in Lefort *et al*. [10]). This method requires to provide a list of targeted metabolites for which the quantification has to be performed. This list is naturally provided by the UPLC dosages in the piglet plasma dataset, but no such natural choice is available for the simulated dataset.

The quality of the quantification was assessed as already described in Section “Evaluation of the joint alignment”, by correlation between estimated quantifications and simulated ones (simulated dataset) or either by correlation between estimated quantifications and UPLC concentrations (piglet plasma dataset). Note that **rDolphin** produces a quantification for several regions of interest that it has identified in the metabolite pure spectrum. We chose to keep only the highest correlation with the UPLC concentrations in our final results in order to show the “best case scenario” of **rDolphin**.

## 4 Results and discussion

### 4.1 Evaluation of ASICS joint alignment procedure

Fig. 4 provides cosine similarities between initial and processed pure spectra for the simulated dataset. It shows that the joint alignment outperforms the other methods. In addition, differences between methods (*p*-value *<* 0.001; Durbin test) as well as pairwise differences (*p*-values *<* 0.001 for all pairs; Durbin post-hoc test) were all found significant.

**Figure 4:**
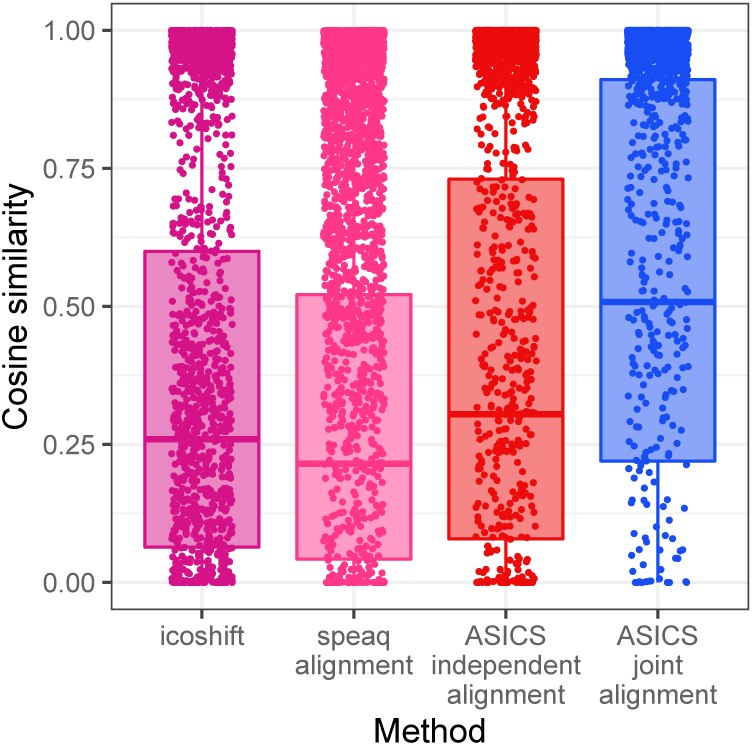
Cosine similarity between the true contribution of **g**_*j*_ to the simulation of **f**_*i*_ (ground truth) and the result of the alignment of **g**_*j*_ on **f**_*i*\*_. Alignments were performed with icoshift, **speaq** or **ASICS** (independent and joint versions) for 100 reference spectra corresponding to the 100 simulations). Points correspond to the cosine similarity of the 30 more concentrated metabolites in every simulation.

The median cosine similarity for **ASICS** joint alignment is equal to 0.51 overall but increases to 0.99 when computed on the 30 more concentrated metabolites only. This is explained by the fact that peaks of very lowly concentrated metabolites are usually under noise signal in the complex spectra and are thus not or poorly detected. For these upmost concentrated metabolites, the median cosine similarity is equal to 0.97 for icoshift and to 0.90 for speaq, both results still significantly differ from **ASICS** joint alignment performances (*p*-values *<* 0.001 in both cases; Durbin post-hoc test).

A similar positive impact of the joint alignment was also obtained on subsequent quantifications for the simulated dataset (Fig. S1 and Table S2 of the Supplementary material). More precisely, the results showed that, even if the sensitivity across methods is not significantly different (*p*-value = 0.60; Kruskal-Wallis test), the specificity and also the correlation between simulated and estimated quantifications were improved by **ASICS** joint alignment (*p*-values *<* 0.001 for both; Kruskal-Wallis and Durbin tests respectively). Median correlations were equal to 0.30 for **ASICS** independent alignment, to 0.32 for icoshift alignment, to 0.34 for speaq alignment and to 0.35 for **ASICS** joint alignment (*p*-values *<* 0.001; Durbin post-hoc tests). Again, median correlation of **ASICS** joint alignment increased to 0.79 when considering only the 30 upmost concentrated metabolites (between 50% and 60% of estimated quantifications were equal to 0).

The evaluation of the impact of the alignment on the quality of the quantification for the piglet plasma dataset exhibited a similar trend (Fig. S2 and Table S3 of the Supplementary material). Correlations between quantifications and UPLC dosages were found higher with **ASICS** joint alignment (median = 0.61) than with **ASICS** independent alignment (median = 0.44; *p*-value = 0.08; Durbin post-hoc test), **speaq** alignment (median = 0.21; *p*-value *<* 0.001, Durbin post-hoc test) or icoshift alignment (median = 0.32; *p*-value *<* 0.001, Durbin post-hoc test). In particular, **ASICS** joint alignment allows to improve the quality of alignment and subsequent quantification of metabolites for which the pure spectrum has a small number of peaks. For instance, the glycine has a pure spectrum with only one peak. In the complex spectra on Figure 5, the actual peak of glycine is at 3.57 ppm. However, with independent alignment, pure spectra of glycine were usually aligned around 3.56 ppm (red spectra). Thus, the correlation between UPLC concentrations and estimated quantifications was equal to 0.07 instead of 0.88 with a joint alignment.

**Figure 5:**
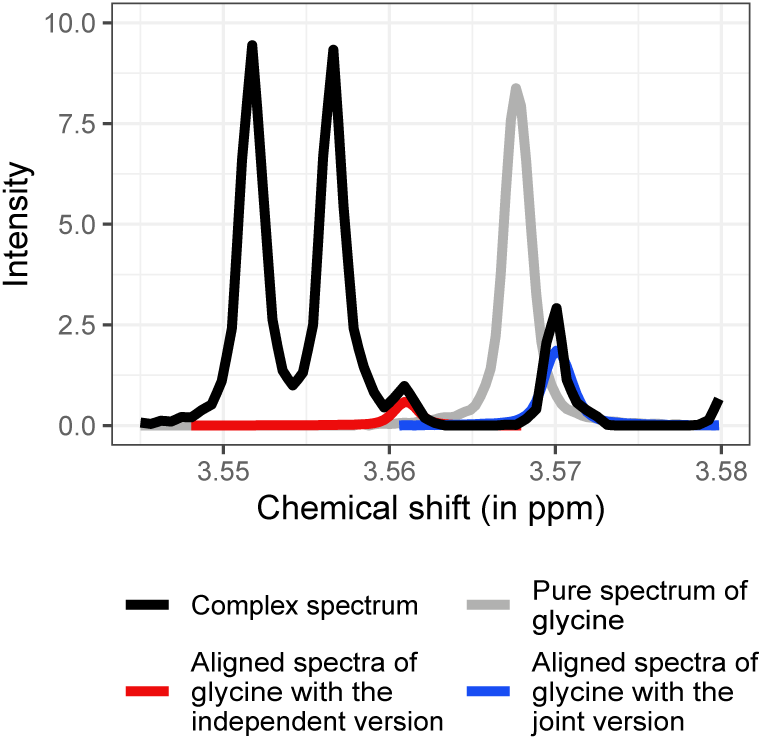
Glycine pure spectrum aligned on every complex mixture spectrum by **ASICS** independent or joint alignments.

### 4.2 Evaluation of ASICS joint quantification

On the simulated dataset, the quality of metabolite identification was found to be opposite for sensitivity and specificity. The best method in terms of sensitivity was **ASICS** joint quantification with a single cleaning step and the worst methods were **ASICS** independent quantification and **ASICS** joint quantification with *r*_*c*_ = 50%, both being very stringent on the identified metabolites (Fig. 6a). On the contrary, these latter two methods were the ones with the best specificity, whereas **ASICS** joint quantification with a single cleaning step achieved the worst specificity (Fig. 6b).

**Figure 6:**
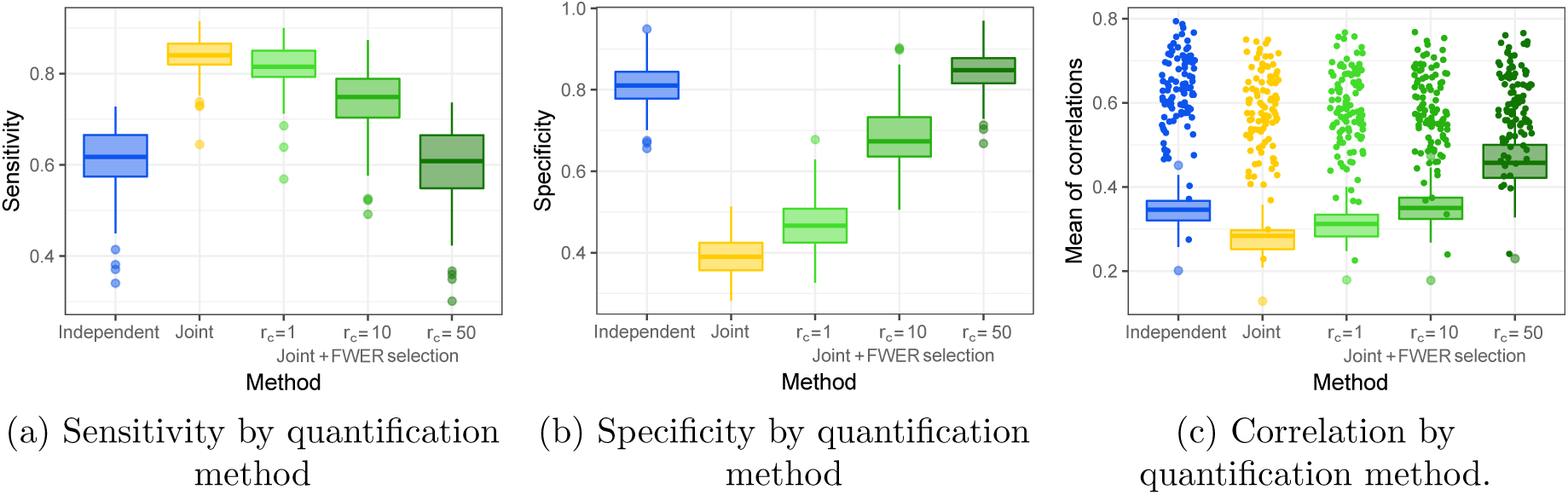
Comparison of quantification methods based on three indicators. Points on Figure 6c correspond to correlations of the 30 most concentrated metabolites.

From the quantification point of view (Fig. 6c), **ASICS** joint quantification with *r*_*c*_ = 50% is the method that achieves the best performances (median correlation equal to 0.46, whereas all the others are below 0.4; *p*-value *<* 0.001 for each pairwise comparison; Durbin post-hoc tests). When looking at the two methods with the highest specificity (**ASICS** independent quantification and **ASICS** joint quantification with *r*_*c*_ = 50%), the quantification was found better with the joint approach (median correlation equal to 0.35 and 0.46 respectively; *p*-value *<* 0.001; Durbin post-hoc test). Indeed, the FWER selection procedure used in **ASICS** independent quantification leads to an under-efficient selection procedure that sets some quantifications to 0 when the joint quantification is able to better estimate their small values (Fig. S3 of the Supplementary material).

Correlations between quantifications and UPLC dosages for the piglet plasma dataset are displayed in Fig. 7 for the different methods. Overall, it shows that **ASICS** joint quantification with *r*_*c*_ = 50% again performs the best on this dataset and gives results significantly better than **rDolphin** (median correlations equal to 0.87 and 0.75, respectively; *p*-value *<* 0.001; Durbin post-hoc test). **rDolphin** performed worse despite the fact that the method was given the metabolites of interest in contrast to **ASICS** that performs its own metabolite identification.

**Figure 7:**
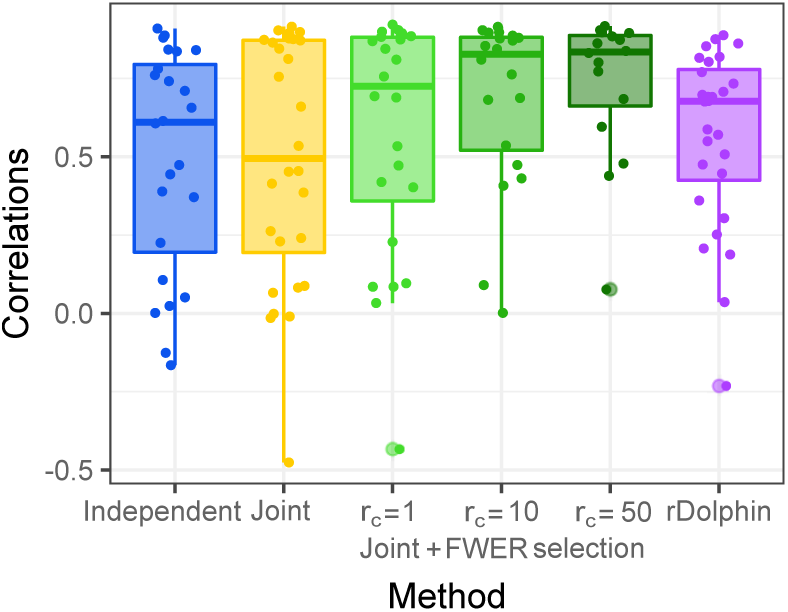
Correlation between quantifications and UPLC dosages by quantification method. Points correspond to all correlations.

In this dataset, amino acid dosages allow to explore a wide variety of concentration values from very concentrated metabolites (alanine or glycine, more than 500*µ*mol/L on average) to very lowly concentrated metabolites (methionine or ornithine, less than 50*µ*mol/L on average). **ASICS** joint quantification allows to address one of the limits of **ASICS** independent quantification described in Lefort *et al*. [10], where quantifications of lowly concentrated metabolites were found of poorer quality. Here, the median correlation of lowly concentrated metabolites (*<* 100*µ*mol/L) was improved by the joint approach with *r*_*c*_ = 50%: median correlations were equal to 0.77 versus 0.50 (*p*-value *<* 0.001; Durbin post-hoc test) for the same two methods (see also examples on the serine and the methionine in Fig. 8).

**Figure 8:**
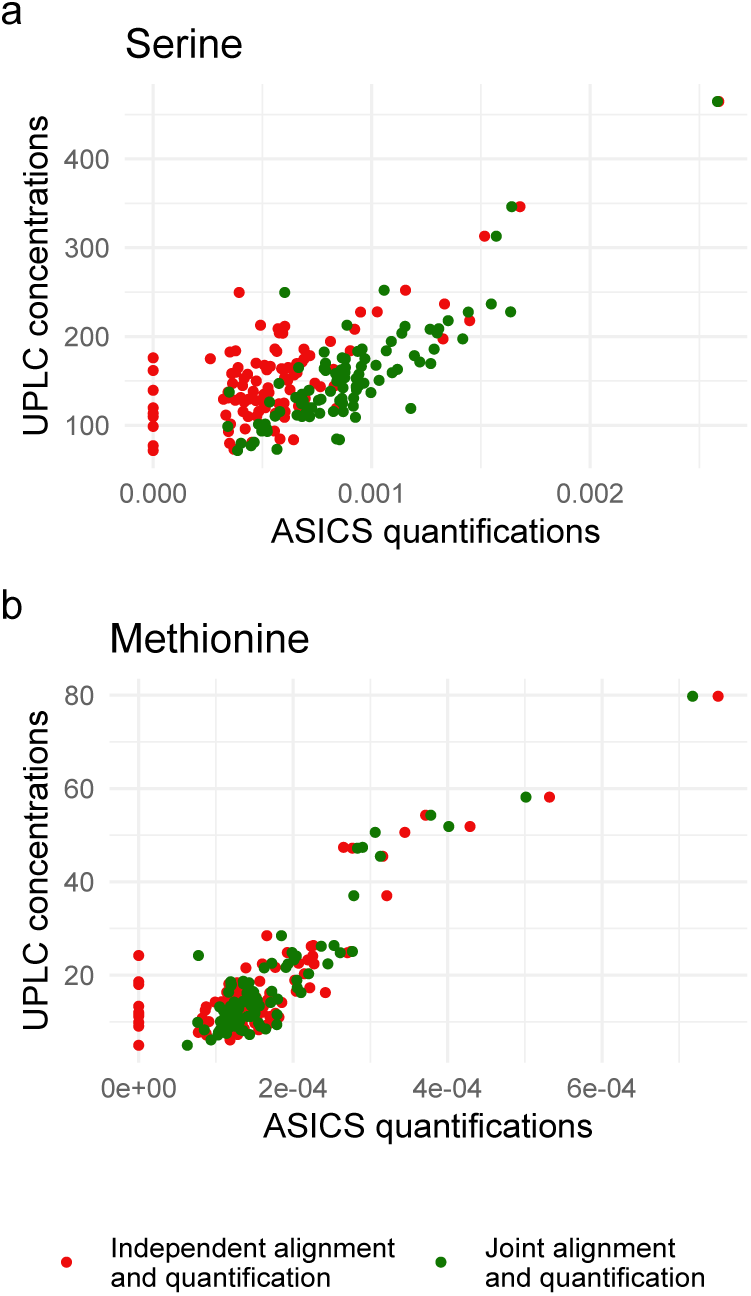
Correlation between quantification and UPLC dosages for (a) serine (155*µ*mol/L on average) and (b) methionine (18*µ*mol/L on average) with independent quantification (red) or joint quantification with *r*_*c*_ = 50% (dark green).

In addition to these two examples, **ASICS** joint quantification also allows to more accurately quantify other types of metabolites that were not identified or were identified only in a few spectra with the FWER selection of **ASICS** independent quantification.

Another case where **ASICS** joint quantification with *r*_*c*_ = 10% provides better results than **ASICS** independent quantification is the case where the pure spectrum of a metabolite has several peaks close to the noise level due to a large number of peaks in this spectrum. This is the case of the lysine, for instance, which has a correlation equal to 0.88 with **ASICS** joint quantification (*r*_*c*_ = 10%) and to 0.42 with **ASICS** independent quantification.

## 5 Conclusion

To the best of our knowledge, **ASICS** joint alignment and quantification approaches are the only automatic approaches that allow to account for multiple samples for automatic identification quantification of metabolites in complex mixture spectra. Both joint steps lead to improved quantification accuracy and a better identification of metabolites present in the complex mixture. In particular, the joint approaches are efficient to help identify metabolites with low concentrations, which are hard to distinguish from the noise level. This is true even when using the joint approaches in combination with stringent pre-filtering steps (*r*_*c*_ = 50%), which are necessary to control the number of false identifications. Finally, with the flexibility offered by the setting of a less stringent pre-filtering step (*r*_*c*_ = 1% or *r*_*c*_ = 10%), the user can also quantify very lowly concentrated targeted metabolites that are known to be present in the complex mixture. Overall, the joint approaches allow to leverage the initial weakness of **ASICS** independent quantification as well as those of most automatic identification methods on the poor identification and quantification of lowly concentrated metabolites.

## 6 Acknowledgement

The authors are grateful to the INRAE metaprogram fund “GISA” (Integrated management of animal health) for the funding of the SuBPig project (Enhancing survival at birth). The PhD fellowship of Gaelle Lefort is supported by the Digital Agriculture Convergence Lab (#DigitAg, http://www.hdigitag.fr/, ANR-16-CONV-0004), by INRAE Mathematics, Computer and Data Sciences, Digital Technologies Division, by INRAE Animal Genetics Division and by INRAE Animal Health Divison. The funders had no role in the study design, analyses, results, interpretation and decision to publish.

The authors are very grateful to the staff of experimental pig facilities (INRAE, 2018. Pig phenotyping and Innovative breeding facility, doi:10.15454/1.5572415481185847E12). They also thank all the participants from GenPhySE, PEGASE and GenESI laboratories (INRAE) and MetaToul-AXIOM plateform for sample and data collection especially Nadine Mézière (INRAE PEGASE) for performing all UPLC dosages, Hélène Quesnel (INRAE PEGASE) for useful discussions and comments and, Roselyne Gautier (MetaToul-AXIOM plateform) for her help in aquisition of NMR spectra.

The authors are grateful to the Genotoul bioinformatics platform Toulouse Occitanie (Bioinfo Genotoul, doi:10.15454/1.5572369328961167E12) for providing computing resources.

The authors also thank Isabelle Pinard for English proofreading and correction.

