## Supplementary Material for "Joint automatic metabolite identification and quantification of a set of ^1^H NMR spectra"

### Supplementary material of the article “Joint automatic metabolite identification and quantification of a set of $^1$H NMR spectra”

Gaëlle Lefort[1,2], Laurence Liaubet[2], Nathalie Marty-Gasset[2], Cécile Canlet[3,4], Nathalie Vialaneix[1,+], Rémi Servien[5,6,+]

[1] *INRAE, UR875 Mathématiques et Informatique Appliquées Toulouse, F-31326 Castanet-Tolosan, France*
[2] *GenPhySE, Université de Toulouse, INRAE, ENVT, F-31326, Castanet-Tolosan, France*
[3] *INRAE, Université de Toulouse, ENVT, Toxalim, 31027 Toulouse, France*
[4] *Axiom Platform, MetaToul-MetaboHUB, National Infrastructure for Metabolomics and Fluxomics, Toulouse, France*
[5] *INRAE, Univ. Montpellier, LBE, 102 Avenue des étangs, F-11000 Narbonne, France*
[6] *INTHERES, Université de Toulouse, INRAE, ENVT, Toulouse, France*
[+] *these authors contributed equally to this work*
{*gaelle.lefort,remi.servien*}*@inrae.fr*

**S4 Comparison of quantification methods** **7**

### S1 Common preprocessing step

**Algorithm S1** Preparation of a common aligned library.

**Require:** user defined ratio of evidence, $r_c \in ]0, 1]$

**for all** $j = 1, \ldots, p$ **do** ▷ Cleaning step 1
    **for all** $i = 1, \ldots, n$ **do**
        Perform independent cleaning steps (based on the presence of peaks of $\mathbf{g}_j$ in $\mathbf{f}_i$) **return** kept metabolites for $\mathbf{f}_i$, $\mathcal{S}_i$
    **end for** ▷ End of Cleaning step 1
    Metabolites $j$ used to fit model (4) are the ones such that: $\frac{|\{j \in \mathcal{S}_i,\ i=1,\ldots,n\}|}{n} \geq r_c$
**end for**
**for all** $i = 1, \ldots, n$ **do** ▷ Cleaning step 2 (optional)
    Perform alignment of the reference library and quantification of $\mathbf{f}_i$ and FWER selection **return** selected metabolites for $\mathbf{f}_i$, $\mathcal{S}'_i$
**end for**
**for all** $j = 1, \ldots, p$ **do**
    Metabolites $j$ used to fit model (4) are the ones such that: $\frac{|\{j \in \mathcal{S}'_i,\ i=1,\ldots,n\}|}{n} \geq r_c$
**end for** ▷ End of Cleaning step 2
**for all** $j = 1, \ldots, p$ **do** ▷ Global alignment
    Perform a joint alignment as described in Section "Joint alignment of the reference library" **return** global shifts $(s_{ij})_{i=1,\ldots,n}$
    Align $\mathbf{g}_j$ using the global shift $\tilde{s}_j = \text{median}(s_{ij})_{i=1,\ldots,n}$
**end for** ▷ End of Global alignment
**for all** $j = 1, \ldots, p$ **do** ▷ Local alignment
    Perform local alignment of $\mathbf{g}_j$ on a reference complex spectrum $\mathbf{f}^{\text{ref}}$ defined as

$$\mathbf{f}^{\text{ref}} = \underset{i=1,\ldots,n}{\arg\max} \frac{1}{n} \sum_{i'=1}^{n} \text{Cor}_{\text{FFT}}(\mathbf{f}_i, \mathbf{f}_{i'}).$$

**end for** ▷ End of Local alignment
**return** Common aligned reference library **G**

### S2 Experimental data for the evaluation

#### S2.1 Simulated spectra

To assess the performances of joint alignment and joint quantification, we first simulated $n$ spectra $(\mathbf{f}_i)_{i=1,\ldots,n}$ with metabolites in known concentrations, $\tilde{b}_{ij}$, from some of the $p$ pure spectra $(\mathbf{g}_j)_{j=1,\ldots,p}$ present in **ASICS** reference library. Five steps were necessary to simulate spectra:

1. a common set of metabolites was selected from the $p$ pure spectra by using $p$ independent Bernoulli random variables with parameter $r = 1/2$;

2. to introduce individual variations between the $n$ simulated complex spectra, $d = 2$ additional metabolites were randomly chosen among all the metabolites, independently for each simulated complex spectra. More precisely, if the metabolite was already present in the common set of selected metabolites (respectively absent), it was removed (respectively added) in the set of selected metabolites for this specific complex spectrum. For $i = 1, \dots, n$, this led to a maximum of four different metabolites between any two complex mixture spectra. In addition, we will denote $p_i$ the number of metabolites present in the $i$th complex mixture spectrum;

3. $\forall\, i = 1, \dots, n$ and $j = 1, \dots, p_i$, ground truth quantifications, $(\tilde{\mathbf{b}}_{ij})_j = (\tilde{\boldsymbol{\beta}}_{ij})_j/(u_j)_j$, were then simulated using $p_i$ independent normal distributions $\mathcal{N}(\mu_1, \sigma_1 = 0.3\mu_1)$ where $\mu_1$ was itself generated from a log-normal distribution of parameters $\mu_2 = -8$ and $\sigma_2 = 2$. Quantifications smaller than 0 were set to 0, as well as quantifications larger than 1 that were set to 1, to avoid an unrealistically large skewness in the simulated quantifications;

4. for each metabolites, $\mathbf{g}_j$ global shifts were simulated independently for each spectra $\mathbf{f}_i$ using negative binomial distributions $s_{ij} \sim NB(2, 0.25)$ and local shifts were simulated independently using normal distributions $\tau_{ijl} \sim \mathcal{N}(0, 0.09)$ with $l$ corresponding to the $l$th peak of the pure spectrum $\mathbf{g}_j$ in the complex spectrum $\mathbf{f}_i$. The final overall shift for this peak was then obtained as $r_{ijl} = \min(s_{ij} + \tau_{ijl}, m_1)$ with $m_1 = 0.02$. Finally, the direction of the shift (left or right), $\alpha_{ijl}$, was chosen using a Bernoulli distribution of parameter 0.5;

5. the simulated complex spectra $\tilde{\mathbf{f}}_i$ were computed as follows: for all chemical shift $t$,

$$\tilde{\mathbf{f}}_i(t) = \sum_{j=1}^{p_i} \tilde{\mathbf{b}}_{ij} u_j \mathbf{g}_j \left(t + (2\alpha_{ijl(t)} - 1) r_{ijl(t)}\right) \qquad \text{(S1)}$$

   with $l(t)$ the peak at position $t$ (if any), $u_j$ the number of protons of the $j$th metabolite. Then, a noise was added based on Equation (1):

$$\mathbf{f}_i = \epsilon_1 \tilde{\mathbf{f}}_i + \epsilon_2$$

   with $\epsilon_1 \sim \mathcal{N}(0, \omega_1^2 = 0.09)$ and $\epsilon_2 \sim \mathcal{N}(0, \omega_2^2 = 0.07)$.

Finally, the $n$ complex spectra were normalized by the area under the curve.

#### S2.2 Plasma spectra of newborn piglets: experimental protocol

**Ethics statement** This study was conducted in accordance with the French legislation on experimentation and ethics. The French Ministry of Agriculture authorized this experiment on living animals at the INRAE facilities (UE1372 GenESI Génétique, Pig phenotyping and Innovative breeding facility, `doi:10.15454/1.5572415481185847E12`) with the agreement number APAFiS for animal housing and the agreement number #13648-2018020417291866 v4 for the protocol.

**Plasma sample collection** Blood (approximately 5 mL) of the 97 piglets was collected individually on piglets from the umbilical cord and placed in heparinized tubes. Plasma was prepared by low-speed centrifugation (2,000 g for 10 min at 4°C) and stored at −80°C until further analysis.

**NMR protocol** Each sample of plasma (200 $\mu$L) was diluted in 500 $\mu$L phosphate buffer prepared in deuterated water (0.2 M, pH 7.0) containing TSP (1.17 mM) as internal standard, vortexed, centrifuged at 5000 g for 15 min at 4°C, and 600 $\mu$L transferred into 5 mm NMR tube. All $^1$H NMR spectra were acquired on a Bruker Avance III HD NMR spectrometer (Bruker Biospin, Rheinstetten, Germany) operating at 600.13 MHz for $^1$H resonance frequency and at 300K, using the Carr-Purcell-Meiboom-Gill (CPMG) spin-echo pulse sequence. Spectrum preprocessing (group delay correction, solvent suppression, apodization, fourier transformation, zero order phase correction, internal referencing, baseline correction and window selection) was perform using the R package **PepsNMR** (version 1.2.1) with the TSP peak for internal reference. Finally, all spectra were aligned with each other using the method implemented in the **ASICS** package.

**HPLC protocol** Plasma amino acid concentrations were obtained using an ultra HPLC system (Waters Acquity Ultra Performance LC system, Waters, Guyancourt, France) coupled to an Acquity tunable UV detector and a mass detector (SQD detector) to identify the few coeluting chromatographic peaks. The column was a MassTrak AAA column (2.1 × 150 mm). Amino acid derivatization was performed with using an AccQ·Tag Ultra derivatization (MassTrak AAA Waters, Milford, MA). Norvaline was used as internal standard. The Empower 2chromatography software (Waters corporation, Milford, MA, USA) was used for instrument control and data acquisition.

Table S1. Minimum, maximum and median concentrations for each metabolites dosed with UPLC.

| Concentrations (in $\mu$mol/L) | Minimum | Maximum | Median | Concentrations (in $\mu$mol/L) | Minimum | Maximum | Median |
|---|---|---|---|---|---|---|---|
| 3-Methylhistidine | 3.16 | 19.22 | 7.58 | Isoleucine | 10.71 | 123.22 | 46.89 |
| Alanine | 270.08 | 1939.22 | 855.93 | Leucine | 24.09 | 229.82 | 79.58 |
| Arginine | 25.49 | 150.97 | 69.35 | Lysine | 76.75 | 388.93 | 218.82 |
| Asparagine | 16.57 | 130.55 | 49.66 | Methionine | 4.97 | 79.77 | 13.43 |
| Aspartic Acid | 1.68 | 57.61 | 10.02 | Ornithine | 11.32 | 70.94 | 34.00 |
| Carnosine | 1.11 | 25.02 | 14.23 | Phenylalanine | 9.00 | 107.22 | 55.40 |
| Citrulline | 39.99 | 152.44 | 80.23 | Proline | 86.85 | 384.19 | 169.51 |
| Cysteine | 10.98 | 42.38 | 22.75 | Sarcosine | 1.78 | 63.20 | 18.49 |
| Ethanolamine | 12.61 | 78.24 | 27.68 | Serine | 71.61 | 464.62 | 147.19 |
| Glutamine | 97.29 | 663.70 | 303.11 | Taurine | 19.93 | 214.85 | 59.22 |
| Glutamic Acid | 44.34 | 567.18 | 153.17 | Threonine | 77.81 | 262.25 | 141.75 |
| Glycine | 177.70 | 1902.22 | 473.74 | Tryptophan | 11.97 | 26.83 | 19.30 |
| Histidine | 24.98 | 256.10 | 96.92 | Tyrosine | 19.13 | 171.25 | 54.26 |
| Hydroxyproline | 42.56 | 140.38 | 70.05 | Valine | 161.72 | 424.20 | 291.98 |

### S3 Comparison of alignment methods

#### S3.1 Simulated dataset

The rate of null quantification is computed on the metabolites identified in at least one complex mixture. It is given by the following formula

$$\text{Rate of null quantification} = 1 - \frac{\sum_{i=1}^{n} \sum_{j=1}^{p} \mathbf{1}_{\{\beta_{ij}>0\}}}{n \sum_{j=1}^{p} \mathbf{1}_{\left\{\sum_{i=1}^{n} \beta_{ij}>0\right\}}}$$

(average frequency of unidentification for metabolites that have been identified at least once). In particular, the rate of null quantification is low if the identified (resp. unidentified) metabolites are identified (resp. unidentified) in all complex spectra, *i.e.*, if the identification are consistent accross complex spectra.

| | **icoshift** | **speaq** | **independent** |
|---|---|---|---|
| **speaq** | 1.00 | - | - |
| **independent** | 0.95 | 0.91 | - |
| **joint** | 0.84 | 0.90 | 0.52 |

(a) Sensitivity (global $p$-value = 0.60; Kruskal-Wallis test)

| | **icoshift** | **speaq** | **independent** |
|---|---|---|---|
| **speaq** | 0.99 | - | - |
| **independent** | 0.005 | 0.003 | - |
| **joint** | 0.04 | 0.05 | $< 0.001$ |

(b) Specificity (global $p$-value $< 0.001$; Kruskal-Wallis test)

| | **icoshift** | **speaq** | **independent** |
|---|---|---|---|
| **speaq** | 0.41 | - | - |
| **independent** | $< 0.001$ | $< 0.001$ | - |
| **joint** | 0.005 | $< 0.001$ | 0.83 |

(c) Null quantification rate (global $p$-value $< 0.001$; Kruskal-Wallis test)

| | **icoshift** | **speaq** | **independent** |
|---|---|---|---|
| **speaq** | $< 0.001$ | - | - |
| **independent** | $< 0.001$ | $< 0.001$ | - |
| **joint** | $< 0.001$ | $< 0.001$ | $< 0.001$ |

(d) Correlation between simulated and quantified metabolites (global $p$-value $< 0.001$; Durbin test)

Table S2. $p$-values of post-hoc Nemenyi tests for sensitivity, specificity and null quantification rate or Durbin tests for correlation for the comparison between each pair of alignment methods (icoshift, **speaq**, **ASICS** independent and joint alignment). **ASICS** independent quantification was performed after library alignment for all methods.

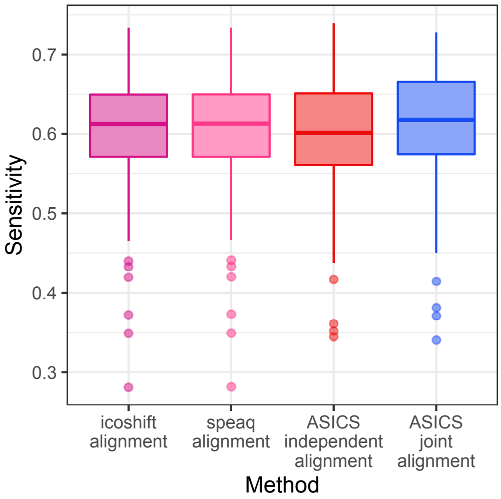

**(a)** Sensitivity by alignment method

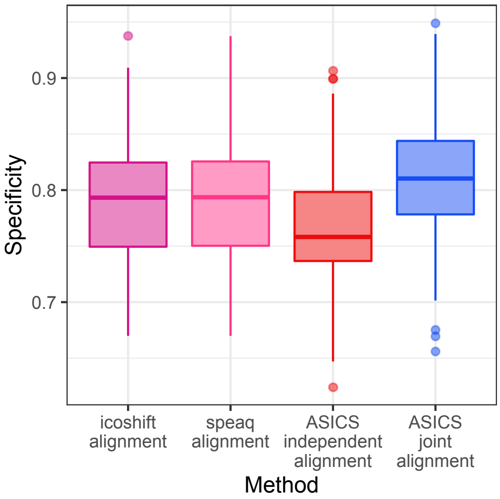

**(b)** Specificity by alignment method

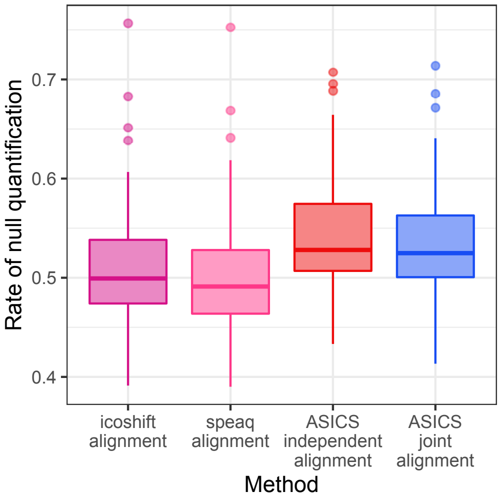

**(c)** Null quantification rate by alignment method

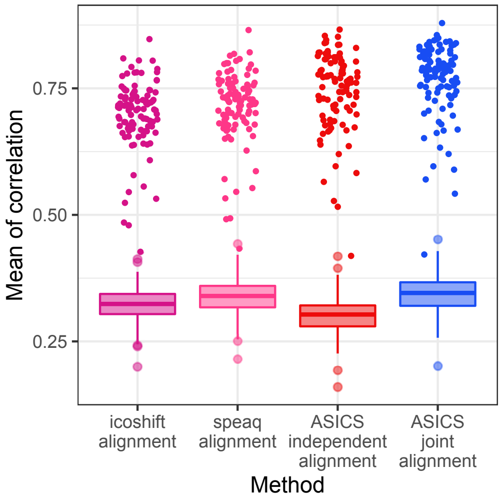

**(d)** Correlation between simulated and quantified metabolites by alignment method.

**Fig. S1.** Comparison of alignment methods based on four indicators. Points on Figure S1d correspond to the correlation obtained for the 30 most concentrated metabolites. **ASICS** independent quantification was performed after library alignment for all methods.

#### S3.2 Piglet plasma dataset

| | **icoshift** | **speaq** | **independent** |
|---|---|---|---|
| **speaq** | 0.71 | - | - |
| **independent** | 0.003 | 0.007 | - |
| **joint** | $< 0.001$ | $< 0.001$ | 0.08 |

Table S3. $p$-values of Durbin post-hoc tests for correlations between quantifications and UPLC dosages between each pair of alignment methods (global $p$-value $< 0.001$; Durbin test). **ASICS** independent quantification was performed after library alignment for all methods.

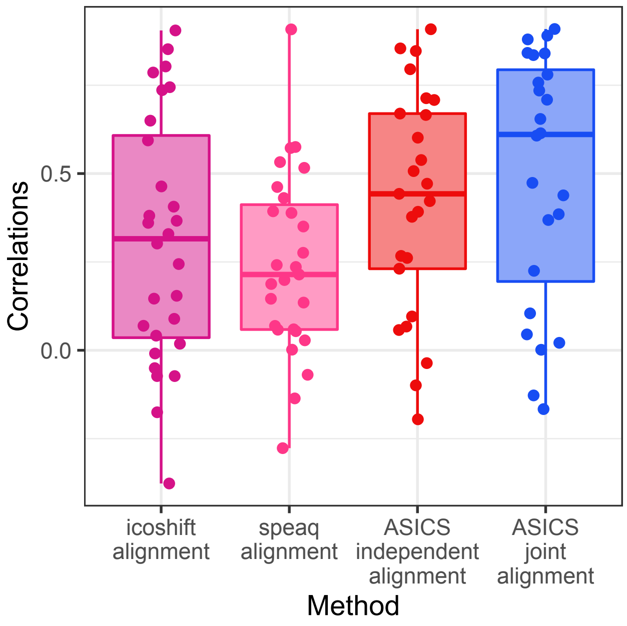

**Fig. S2.** Correlations between quantifications and UPLC dosages using three different alignment methods. **ASICS** independent quantification was performed after library alignment for all methods. Points correspond to every individual correlations.

### S4 Comparison of quantification methods

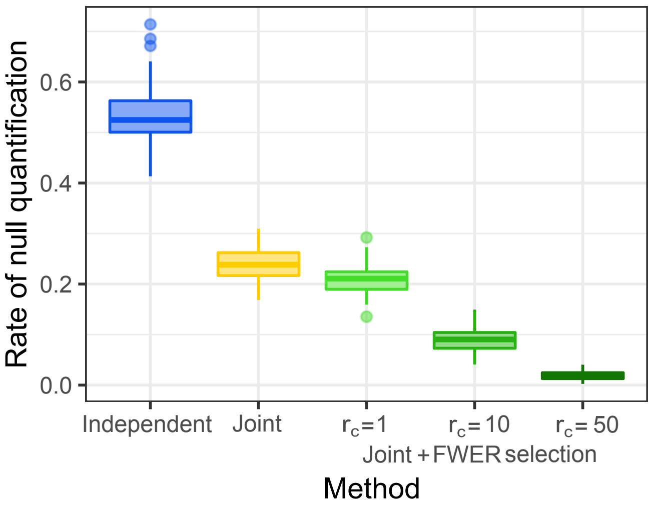

**Fig. S3.** Null quantification rate by quantification method.
